# A cellular automaton for modeling non-trivial biomembrane ruptures

**DOI:** 10.1101/429548

**Authors:** Abhay Gupta, Ganna Reint, Irep Gözen, Michael Taylor

## Abstract

A novel cellular automaton (CA) for simulating biological membrane rupture is proposed. Constructed via simple rules governing deformation, tension, and fracture, the CA incorporates ideas from standard percolation models and bond-based fracture methods. The model is demonstrated by comparing simulations with experimental results of a double bilayer lipid membrane expanding on a solid substrate. Results indicate that the CA can capture non-trivial rupture morphologies such as floral patterns and the saltatory dynamics of fractal avalanches observed in experiments. Moreover, the CA provides insight into the poorly understood role of inter-layer adhesion, supporting the hypothesis that the density of adhesion sites governs rupture morphology.

## Introduction

Under mechanical stress, biological membranes have been shown to display non-trivial pore morphologies and dynamics^1^ in addition to the previously known circular transient pores. The peculiar morphologies comprise large flower-like shapes growing continuously referred to as ‘floral’ pores, and fractures appearing intermittently in ‘fractal’ patterns. Both rupture types occur in the exact same experimental conditions with the identical membrane compositions, solid substrates and ambient buffers. The factors which influence the fate of the rupture mode are not exactly understood and the detailed mechanisms remain to be elucidated.

We hypothesize that the density of adhesion sites of the membrane to the underlying layer, defines the rupture type. In our experiments, where a double lipid bilayer membrane (DLBM) is formed on a solid substrate via self-spreading of a lipid reservoir, the adhesion is most likely a result of pinning between the two bilayers mediated by Ca^2+1, 2^. In a recently reported experimental system where fractal pores in solid-supported membranes were observed, the adhesion was created via biotin-avidin bridging of the two bilayers^3^. The water layer between the two bilayers in a DLBM stack through which the Ca^2+^-mediated pinning is established, is maximum a few tens of nanometers thick. An intact DLBM with a diameter of 100 μm and an inter-bilayer space of 10 nm has a volume of about 750 fL^4^. This makes the direct observations, for instance the visualization of the labeled Ca^2+^ ions, challenging. Therefore, we utilize computer modelling, which allows us to independently tune the potentially influential parameters affecting the rupture morphology to understand both their individual impact and, consequentially, the detailed mechanisms involved.

Researchers have noted similarity between the floral and fractal lipid rupture morphologies and the interaction of immiscible fluids in porous media^1, 4^. In particular, lipid fracture is related to viscous fingering and percolation phenomena. These types of instabilities have been of interest to researchers investigating oil recovery^5^, fluid mixing^6^, soil physics^7^, and biological tissue and organ engineering^8, 9^.

One of the most well-established computational paradigms for exploring percolation and growth phenomena is spatial simulation, of which cellular automata (CA) are a prominent example^10, 11^. CA trace their roots back to von Neumann’s desire to create self-replicating machines^12, 13^ and were popularized by Conway’s discovery of the Game of Life^14^. Subsequent academic interest in CA owes much to the work of Wolfram^15^. A cellular automaton consists of a grid of cells (commonly in one or two dimensions) that are assigned a specific state (or states). In the simplest of CA, this state is binary, 0 or 1, on or off, alive or dead, etc.; although, this is not a necessity. These states evolve through the application of a few very simple rules, e.g. “randomly select a 0 neighbor of a 1 cell and make it 1.” Despite their simplicity, CA are able to capture a range of complex behavior, even emergent behavior, in the areas of growth, aggregation, segregation, and percolation^10, 11^.

While CA are relatively simple, numerical methods for modeling fracture typically are not. This is due to the fact that ruptures involve discontinuities that hamper discretizations based on spatial derivatives. Peridynamics is a continuum mechanics theory that avoids this problem by being based on an integral equation of motion^16, 17^. In its mesh-free discretization, it models particles interacting via bonds, which can break causing fracture^18^. Our recently published mesh-free peridynamic model of biomembrane ruptures revealed that the fluid biological membranes which favor fractal morphologies could adopt a non-zero shear modulus^19^. While the model captures circular and floral patterns and their associated dynamics, it does not capture the saltatory dynamics of fractal rupture nor the very fine fractal patterns observed experimentally.

In this work, we propose a new innovative CA incorporating ideas from both standard percolation models as well as our previous peridynamics model that captures circular, floral, and fractal avalanche morphology and their associated dynamics in single framework. We illustrate the CA model through simulations of an expanding DLBM on a solid substrate and compare with experimental results. The goal of this new model is to determine the rules underlying *pattern formation* and *saltatory dynamics* of lipid membrane rupture in order to gain insight on the role of pinning in this process, which is still poorly understood. This is important because a better understanding of pore formation in biological membranes can help in finding possible mechanisms behind cell-integrity related diseases^20–22^ and also would help establish a ground for improvement of related medical treatment methods e.g. drug delivery^23^, gene therapy^24^.

## Materials and Methods

### Preparation of lipid suspension

*Lipids and lipid fluorophore:* Soybean Polar Lipid Extract(SPE); E. coli Polar Lipid Extract(ECPE); 1-oleoyl-2- (6-((4,4-difluoro-1,3-dimethyl-5-(4-methoxyphenyl)-4-bora-3a,4a-diaza-s-indacene-2- propionyl)amino)hexanoyl)-sn-glycero-3- phosphoethanolamine (TopFluor^TM^ TMR PE), were obtained from Avanti Polar Lipids (AL, USA).

*Buffers:* <u>PBS</u>: 5 mM Trizma Base (Sigma Aldrich), 30 mM K_3_PO_4_ (Sigma Aldrich), 30 mM KH2PO4 (Sigma Aldrich), 3 mM MgSO_4_–7H2O (Sigma Aldrich) and 0.5 mM Na_2_EDTA (Sigma Aldrich). The pH was adjusted to 7.4 with H_3_PO_4_. <u>HEPES without CaCl_2_</u> (for rehydration): 10 mM HEPES (Sigma Aldrich), 100 mM NaCl (Sigma Aldrich), the pH was adjusted to 7.8 with NaOH. <u>HEPES with CaCl_2_</u> (ambient buffer for spreading): 10 mM HEPES (Sigma Aldrich), 100 mM NaCl (Sigma Aldrich), 2,5 mM or 4 mM CaCl_2_ (Sigma Aldrich) – accordingly to experimental setup. The pH was adjusted to 7.8 with NaOH.

Lipids and lipid-conjugated fluorophore, all dissolved in chloroform, were mixed in the following ratios: SPE 50 wt %, ECPE 49 wt %, TopFluor TMR PE 1 wt %. The volumes corresponding to the specified mass fractions of lipids, were transferred into a round-bottom flask leading in total to 3000 μg of lipids in 300 μl chloroform (10 mg/ml). The solvent was evaporated in a rotary evaporator at –80 kPa for 6 hours, to form a dry lipid cake at the bottom of the flask. 3 ml of PBS buffer and 30 μl of glycerol were added to the flask and the mixture was placed at +4°C overnight for swelling. The next day the flask was placed in an ultrasonic bath sonicator (VWR) at 30 °C for 15–30 s, to form a lipid suspension containing giant multi- and unilamellar vesicles.

### Surface fabrication

SiO_2_ coatings were produced in the clean room facility MC2 at Chalmers University of Technology, Sweden using MS 150 Sputter system (FHR Anlagenbau GmbH). All depositions were applied on glass cover slips (#1, Menzel-Gläser) which were pre-cleaned with oxygen plasma 2 min at 50 W using Plasma Therm BatchTop PE/RIE m/95. The final film thickness of SiO2 was, for fractal: 10 nm, for mixed patterns: 84 nm and for floral pore formation: 15 nm.

### Spreading

4 μl of the stock lipid suspension described above, was placed on a clean glass cover slip and desiccated for 20 minutes. The resulting dry film was rehydrated with HEPES buffer without CaCl_2_ for 3 minutes. The re-hydrated sample containing lipid vesicles were placed on top of a SiO2 coated glass slides, in HEPES buffer with Ca^2^+ which leads to spreading of the multilamellar lipid vesicles (reservoirs) in form of a double lipid bilayer membrane (Fig. 1).

### Microscopy

A laser scanning confocal microscope (Leica SPX8, Germany) was used to visualize the experiments. A white light laser source was used for excitation of TopFluor^TM^ TMR PE at 544 nm and emission was collected at 560–600 nm by a photomultiplier tube detector.

### Numerical Model

Like the authors’ previous peridynamic study^19^, we consider only the expanding and rupturing distal lipid bilayer in creating a two-dimensional model of the system. Accordingly, our CA are contained within a *n × n* square region of cells. Initially, all cells within a radius *R_i_* are considered to be part of the lipid bilayer. In each generation, the radius of the membrane is increased by one cell, which continues until a final radius *R_f_* = ^*n*^/2 is reached, whereupon the simulation ends. Both *n* and *R_i_* are user-specified parameters.

In addition to a Boolean state governing whether or not a cell is within the lipid membrane, we assign states describing whether or not a cell is a pinning site (Boolean), its tension (floating point number), and whether or not it is ruptured (Boolean). Pinning location and behavior are initialized stochastically, while tension and rupture are governed by fixed stiffness and critical tension parameters, respectively, which are assigned material properties. We describe pinning, tension, and fracture in more detail in the following sections. All simulations were performed using a specially-written program in Matlab 2015b (MathWorks), available in the electronic Supporting Information (SI).

#### Pinning

In the real physical system, pinning between distal and proximal layers is widespread with locations that are unpredictable. To account for this variability in our numerical model, we assign two pseudorandom numbers *θ*_1_,*θ*_2_ ∈ [0,1] to all cells within the lipid. Cells with *θ*_1_ less than a user-defined pinning probability *P_pin_* become a (dilute) pinning site. If, in addition, a pinned cell with *θ_2_* less than a user-defined cluster probability *P_cluster_* become the root of a (dense) pinning cluster.

To create the cluster, we use a *site percolation* model^10^. First, cells in Moore’s neighborhood (i.e., the eight adjacent cells) of the cluster root become a pin with a probability *P_perc_* based on their *θ*_1_ value. Next, cells in Moore’s neighborhood of these new pins also become a pin with a probability *P_perc_* based on their *θ*_1_ value. This process is continued recursively until there are no more neighbors with *θ*_1_ < *P*_perc_, at which point the cluster has reached its terminal size. Fig. 1 shows the schematic representation of the hypothesized dilute and dense pinning as well as a typical CA snapshot with pinning sites in both dilute and dense cases present.

**Figure 1.**
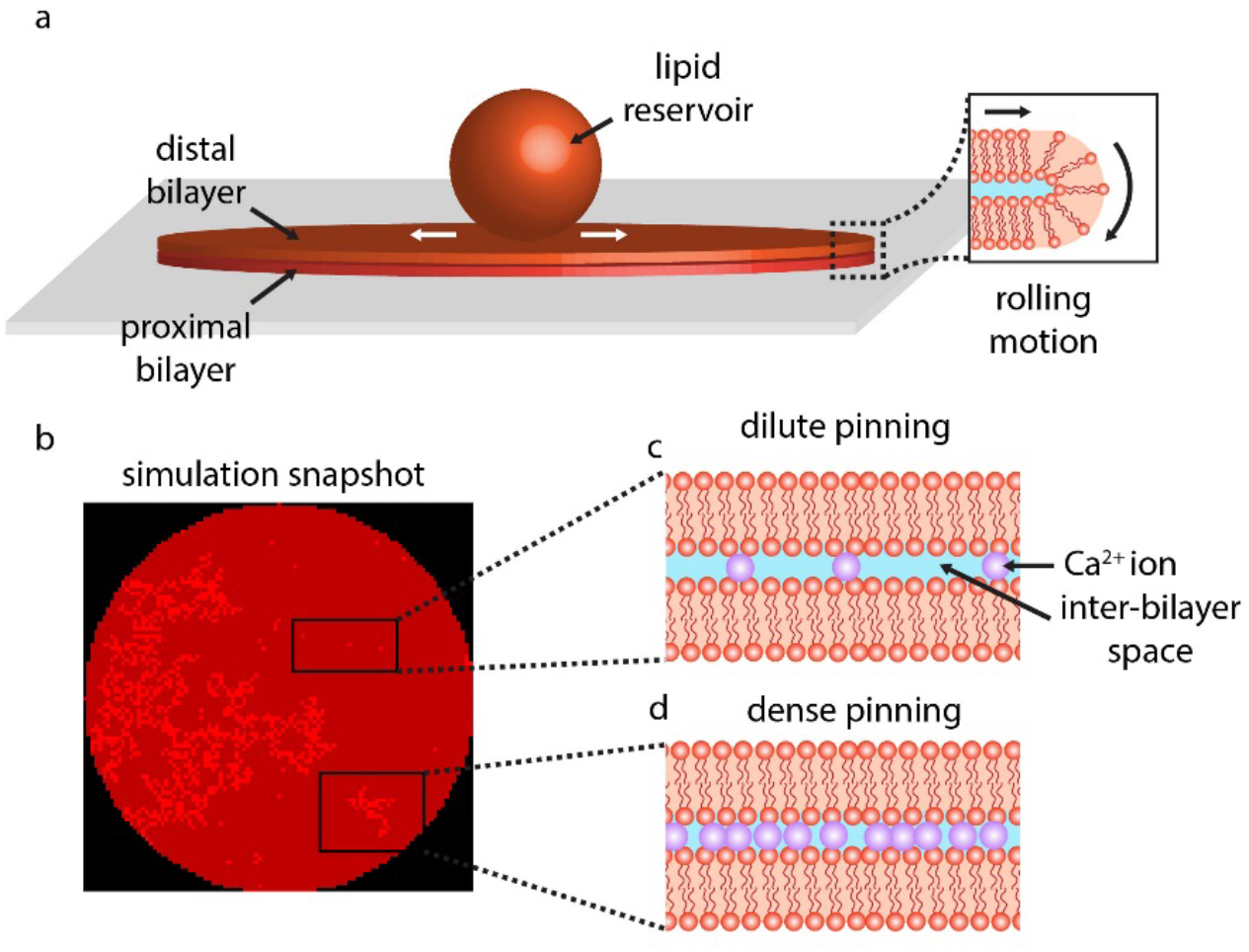
Overview of the experiment and the model. **(a)** The experiment starts with placing a multilamellar reservoir on a SiO_2_ substrate which leads to spreading of the reservoir as a circular double lipid bilayer membrane (DLBM). The peripheries of the circular membrane are depicted open to ease the understanding of the reader. In reality the edges are closed (inset) to avoid the exposure of the hydrophobic moieties of the phospholipid molecules to the aqueous buffer. **(b)** A simulation snapshot showing pinning points (bright red regions). The model is investigating the hypothesis that the type of rupture morphology and associated rupture dynamics, are determined by the density and distribution of pinning. The pinning occurs due to the Ca^2+^ bridging the two bilayers. **(c)** The dilute pinning, relatively larger space in between the pinning sites, cause large pores or floral patterns. **(d)** The dense pinning, where Ca^2+^ ions are packed closely, lead to fractal ruptures. In the simulations, the dilute pinning is programmed by randomly assigning cells within the lipid area as pinned; and dense pinning, by randomly placing pinned clusters based on a site percolation algorithm.

We note that our CA is deterministic. While the system is initialized pseudorandomly, the values of *θ*_1_ and *θ*_2_ as well as the underlying rules remain fixed throughout the simulation.

#### Tension

As the distal lipid layer expands, its tension increases. The presence of pinning sites act as local tension raisers. Accordingly, all pinned cells in our model are assigned a tension *T* comprising both the spreading tension *T_s_* and a pinning tension *T*_p_. In the physical system, the spreading is driven by a gradient between the surface adhesion tension and the internal tension of the multilamellar reservoir (i.e., Marangoni flow). Approximating the membrane as incompressible, the spreading tension at points located distance *r* from the membrane center is^4^ 
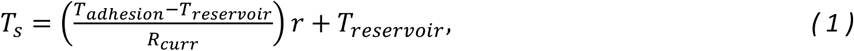
where *T_adhesion_* is the tension due to the lipid’s adhesion to the surface, *T_reservoir_* is the tension of the multilamellar reservoir, and *R_curr_* is the current radius of the lipid membrane. For an expanding membrane, *T_adhesion_* > *T_reservoir_*.

The pinning tension arises from the expanding membrane pulling on the fixed pinned cells. We assume a linear model for the pinning tension governed by a stiffness parameter *c* 
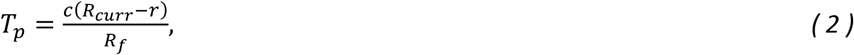
which is higher for those pinned cells closer to the center of the membrane. This is in contrast to the spreading tension, which is higher for those cells nearest outer edge of the membrane. In our model, we choose *T_adhesion_, T_reservoir_*, and *c* such that the total tension *T* = *T_s_* + *T_p_* is highest at the center of the membrane (i.e., the pinning tension dominates) as that gives the best agreement with experiments. Fig. 2a shows typical tension profiles for several *R_curr_* values in a CA with *R_f_* = 100.

**Figure 2.**
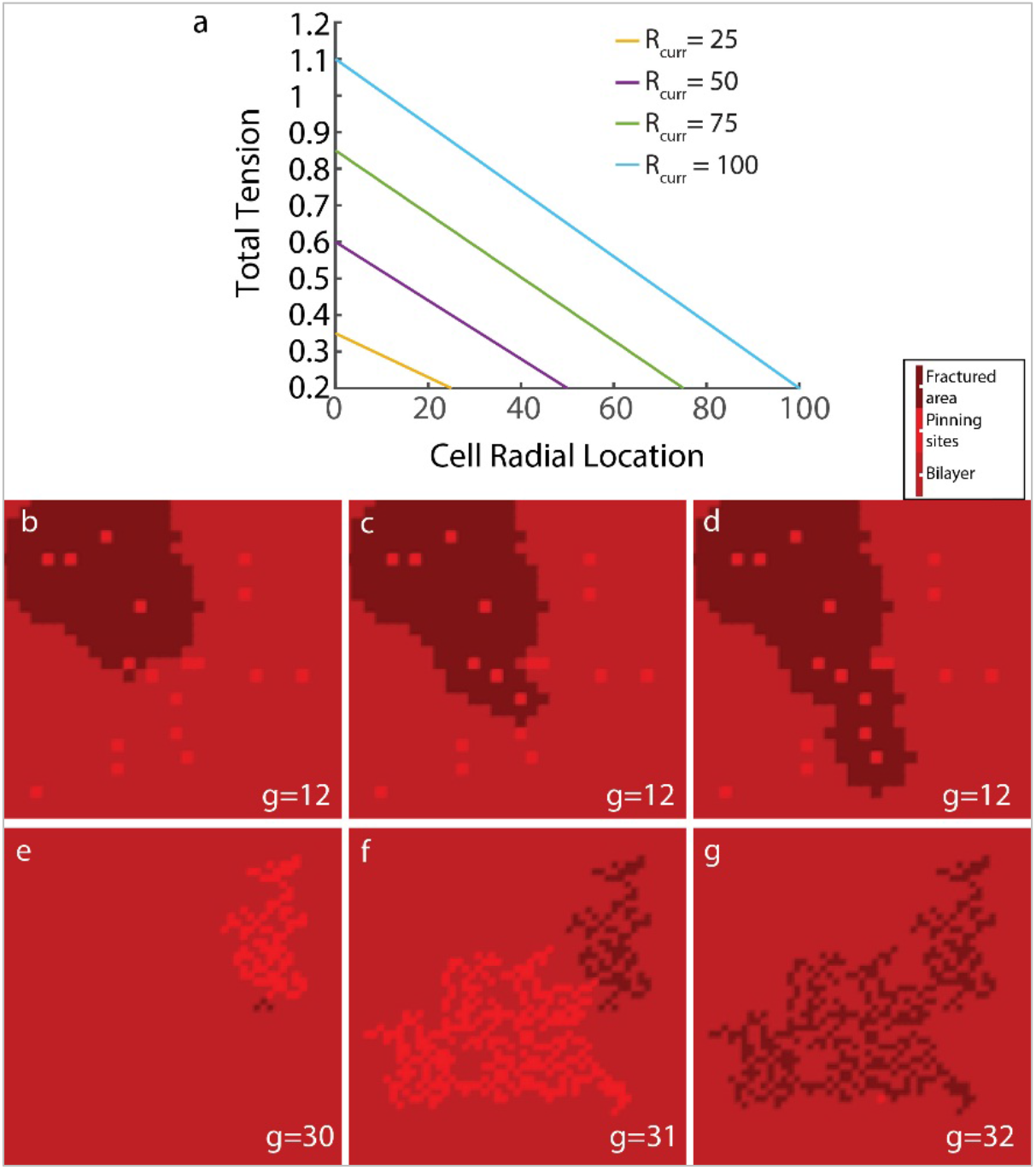
Details of pore formation in the CA model. **(a)** The total tension *T* = *T_s_* + *T_p_* as a function of radius r is shown for various *R_curr_* and a final radius *R_f_* = 100 when the *CTP* = 0. **(b-d)** Typical fracture event for dilute pins. When a floral pore reaches an unbroken pin (b), a new circular pore forms at that pin and grows radially outward. The new pore contacts neighboring unbroken pins, causing new pores to form (c), which grow and cause other neighboring pins to rupture (d). **(e-g)** Typical fracture event for cluster pins. A small broken cluster begets an offshoot cluster of pins (e), which ruptures under higher tension in a succeeding generation, begetting another offshoot cluster (f), which ruptures in turn during a succeeding generation (g).

#### Fracture

In every CA generation, each pinned cell’s tension *T* is checked against a critical tension *T*_crit_. If the tension exceeds the critical tension, the cell is fractured. This straightforward type of fracture criteria is common among bond-based mechanics models, of which our prior peridynamic work is an example^19^. The growth and spread of the rupture are determined by the type of pin the cell is (Fig. 2b-g). For dilute pins (Fig. 2b-d), fracture is an increasing circular pore, the radius of which grows by one cell in each CA generation. Cells within this radius are fractured regardless of their pinning state. If a pore of radius *R_pore_* reaches another pinned cell that hasn’t fractured, it is immediately fractured (regardless of tension). Moreover, all cells within a radius of *R_pore_* of the newly fractured cell are fractured as well. This process of pore growth possibly initiating new pores, is done recursively within a given CA generation and models the continuous linear increase in floral pore area observed experimentally^1^.

For pinned clusters(Fig. 2e-g), rupture growth proceeds differently. Once the critical tension is exceeded by a single cell in a cluster, all cells in the cluster (and any cluster pin cells in adjoining clusters) are fractured simultaneously. Rather than releasing energy by creating a growing circular pore, fractured clusters create a chain of many small static pores. In addition, they generate a new unbroken offshoot cluster with an offshoot probability *P_offshoot_*. This is a realistic assumption since it is anticipated that in the experiments when a pore opens, Ca^2+^ ions in the ambient buffer rapidly penetrates into the interbilayer space, dynamically inducing further pinning. This offshoot cluster is rooted at the neighboring cell with the lowest *θ*_1_ such that *θ*_1_ < *P_offshoot_* and created following the same site percolation process described earlier. If there is no such cell meeting this criterion, no offshoot cluster is formed. The offshoot cluster is initialized with a reduced tension 
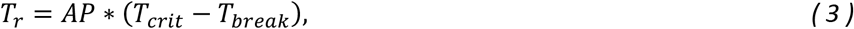
where *T_break_* was the tension that caused the initiating cluster to break and *AP* is a positive avalanche parameter. The reduced tension accounts for the energy released by the fracture of the initiating cluster and subsequent relaxing of the membrane. *AP* governs the saltatory behavior of fractal rupture observed in experiments^1, 3^. When *AP =* 0, saltatory growth is suppressed. As *AP* is increased, the delay between cluster ruptures becomes more pronounced. A more detailed analysis of this behavior is given in the Results and Discussion section.

As noted in the previous section, the tension is defined such that it is highest towards the center of the membrane. Thus, fracture tends to start at those cells close to the center. In our experiments, we observe that fractal ruptures sometimes initiate at the outer edge rather than the center of the membrane. To account for this, we modify the pinning tension (2) for cluster pins based on the number of cells in the cluster *N_cluster_* 
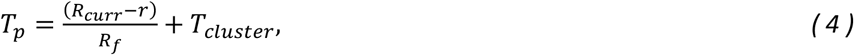
where 
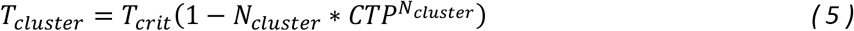
and *CTP ∈* [0,1] is a cluster tension parameter used to differentiate the behavior of small clusters (i.e., those for which *N_cluster_ < 20)* versus larger clusters. For *CTP >* 0.5, *T_cluster_* increases the tension on clusters of all sizes up to a maximum of *T_crit_*; although, its effect is larger on large clusters. In this case, ruptures will tend to start at the outer edge (Fig. 3a). On the other hand, for *CTP* < 0.5, *T_cluster_* reduces the tension on small clusters while increasing the tension of larger clusters. In this case, ruptures tend to initiate towards the center of the lipid (Fig. 3b). Fig. 3c shows *T_cluster_* as a function of cluster size for a range of *CTP* values.

**Figure 3.**
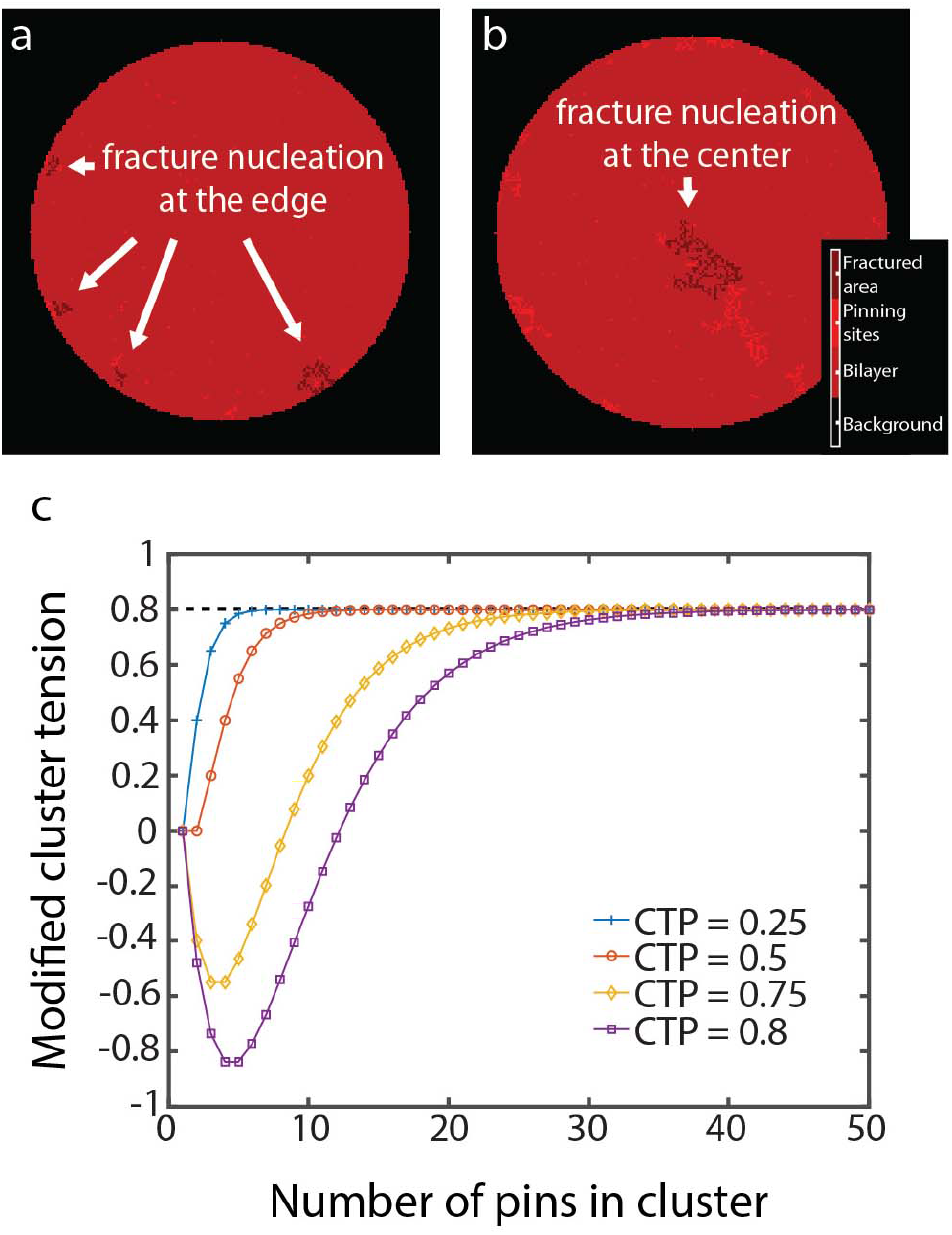
| Effective parameters for location of nucleation. **(a-b)** Simulation snapshots showing the nucleation of fracturing at (a) periphery for non-zero cluster tension parameter (CTP) and **(b)** at the center for CTP = 0 **(c)** The CTP controls the extent to which cluster size affects the modified cluster tension *T*_cluster_, here shown for *T_crit_ =* 0.8.

### Results and Discussion

We start the experiments by placing a lipid reservoir, a multilamellar phospholipid vesicle (MLV), on a SiO2 surface. Upon contact with the surface, the MLV spontaneously starts to wet the surface as a circular double lipid bilayer membrane (DLBM) (Fig. 1a)^1^,^25^). The periphery of the circular membrane in Fig. 1a is depicted open, to facilitate the understanding of the reader. In reality the DBLM is intact, and the edges are closed (inset) to avoid the exposure of the hydrophobic moieties of the phospholipid molecules to the aqueous buffer. The continuous spreading increases the tension of the membrane and when exceeds the lysis tension (5–10 mN m^-1^), the membrane ruptures^1^. The morphology and dynamics of the biomembrane ruptures mainly follow two forms: large floral pores which continuously progress; and fine, fractal pores which occur intermittently, with waiting times in between^19^,^1^). Occasionally, we observe both ruptures on the same membrane patch. In this model, we hypothesize that the number and distribution of the pinning sites, most possibly caused by the Ca^2+^ ion bridging the two bilayers, determine the morphology of the rupture (Fig. 1b-d). Fig. 1b shows a snapshot from our simulations, where pinning sites both individual and in clusters, have been randomly positioned throughout the circular patch (bright red).

The singular pinning points represent individual or very few Ca^2+^ ions (Fig. 1c); and the clusters represent dense regions of Ca^2+^ bridging between the two layers (Fig. 1d).

Fig. 4 shows membranes on a solid substrate and the corresponding CA simulations displaying the two distinct rupture morphologies, floral and fractal, as well as the combination of both (SI Movies). The key parameters used in the CA simulations are given in Table 1. The membrane in the experiments is doped with a fluorescence dye, which makes it visible. The micrographs are taken from top view. In the experiments when a pore opens, the proximal membrane becomes visible through the opening in the distal membrane, represented by half the light intensity^1^. The lipid material which initially resides in the area before the pore opens, now migrates toward the edges of the lipid patch. In the simulations, different colors represent from the most to less intense red: the pinning sites, the bilayer (distal membrane), fractured area (the proximal membrane). The background (surface) is depicted as black.

**Table 1.** Parameters used in the cellular automaton simulations shown in Fig. 4.

|  | Floral | Fractal | Mixed |
| --- | --- | --- | --- |
| $P_{pin}$ | 2% | 0.4% | 4% |
| $P_{cluster}$ | 0% | 100% | 0.08% |
| $P_{perc}$ | NA | *see Eq (6) | 36% |
| $P_{offshoot}$ | NA | 70% | 70% |
| $T_{adhesion}$ | 0.2 | 0.2 | 0.2 |
| $T_{reservoir}$ | 0.1 | 0.1 | 0.1 |
| $T_{crit}$ | 0.8 | 0.75 | 0.8 |
| $T_{cluster}$ | Off | Off | On |
| $C$ | 1 | 1 | 1 |
| $R_i$ | 100 | 100 | 100 |
| $R_f$ | 150 | 150 | 150 |
| $AP$ | NA | 20 | 20 |
| $CTP$ | NA | NA | 0.75 |

The opening and propagation of a floral pore in a circular membrane patch can be observed in Fig 4a in an experiment, and Fig 4b in a simulation where **g** (= *R_curr_* – *R_i_*) represents the CA generation number of the snapshot. The key parameters in the simulation are *P_pin_* = 2% and *P_cluster_* = 0%, meaning that all the pins in the model are of the dilute type and there are no pinned clusters. In experiments, the floral rupture is seen to advance along multiple fronts across membrane, which our CA is able to capture. We note the similarity of the boundary of the rupture front between the simulation and experiment. In general, increasing *P_pin_* (i.e., increasing the number of pin sites) would lead to smaller circular ruptures and a more varied rupture front boundary, whereas decreasing *P_pin_* would lead to larger circular ruptures and a more uniform rupture front boundary. Both cases have been observed experimentally.

Fig 4c and d shows examples of the purely fractal rupture morphology in an experiment and a simulation, respectively. The key difference in the simulation in this case versus the floral case is that we now set *P_cluster_ =* 100%, meaning that all pins belong to a cluster and there are no dilute pins. In order to match the experiments in Fig. 3c, in which ruptures begin toward the center, we turn *T_cluster_* off. In addition, we use a non-constant function for *P_perc_* 
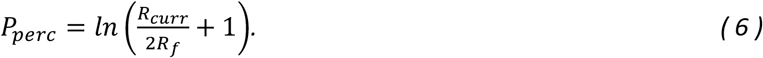

For the parameters used in this study, this leads to 0.288 ≤ *P_perc_* ≤ 0.406. The motivation for (6) is based in part on the observation of some experiments that clusters appear to become larger as the radius increases and also on one of the fundamental concepts of percolation theory-spanning clusters. Spanning clusters are those that fully extend across at least one dimension of a CA. Percolation models are marked by a phase transition at a critical threshold 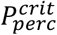, near and beyond which the chances of a spanning cluster appearing increases dramatically^10^. For a two-dimensional system utilizing Moore’s neighborhood, such as the one we use in this study, 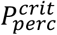 is approximately 0.407^10, 26^, which is just above the upper bound of (6). At 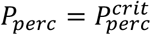, clusters are scale-invariant (i.e., are truly fractal in nature)^10^, and, in 2D, spanning clusters have a fractal dimension of 91/48^27^. In both simulation and experiment, we note that the ruptures begin towards the center of the membrane and grow into fractal structures that nearly reach the outer edge while trapping “islands” of lipid in the interior.

Occasionally, some membrane patches display fractal and floral ruptures simultaneously. Fig 4e shows a typical experiment where the both types of ruptures are visible on the same patch. Fig 4f shows the snapshots from the corresponding CA simulation. We note that the ruptures in Fig. 4e are primarily floral with only 2–3 identifiable clusters near the outer edge that are relatively small. Given the apparent dominance of dilute pinning, we use a very small cluster probability of *P_cluster_* = 0.08%. In addition, since the clusters are relatively small, we use a constant percolation probability sufficiently far from the critical threshold *(P_perc_* = 36%). To account for the clusters initialing closer to the membrane edge, we use a non-zero *T_cluster_* with a *CTP* = 0.75. Comparing experiments with simulation, we see that both initially rupture in a fractal cluster before eventually breaking into a floral pattern, which spreads throughout.

**Figure 4.**
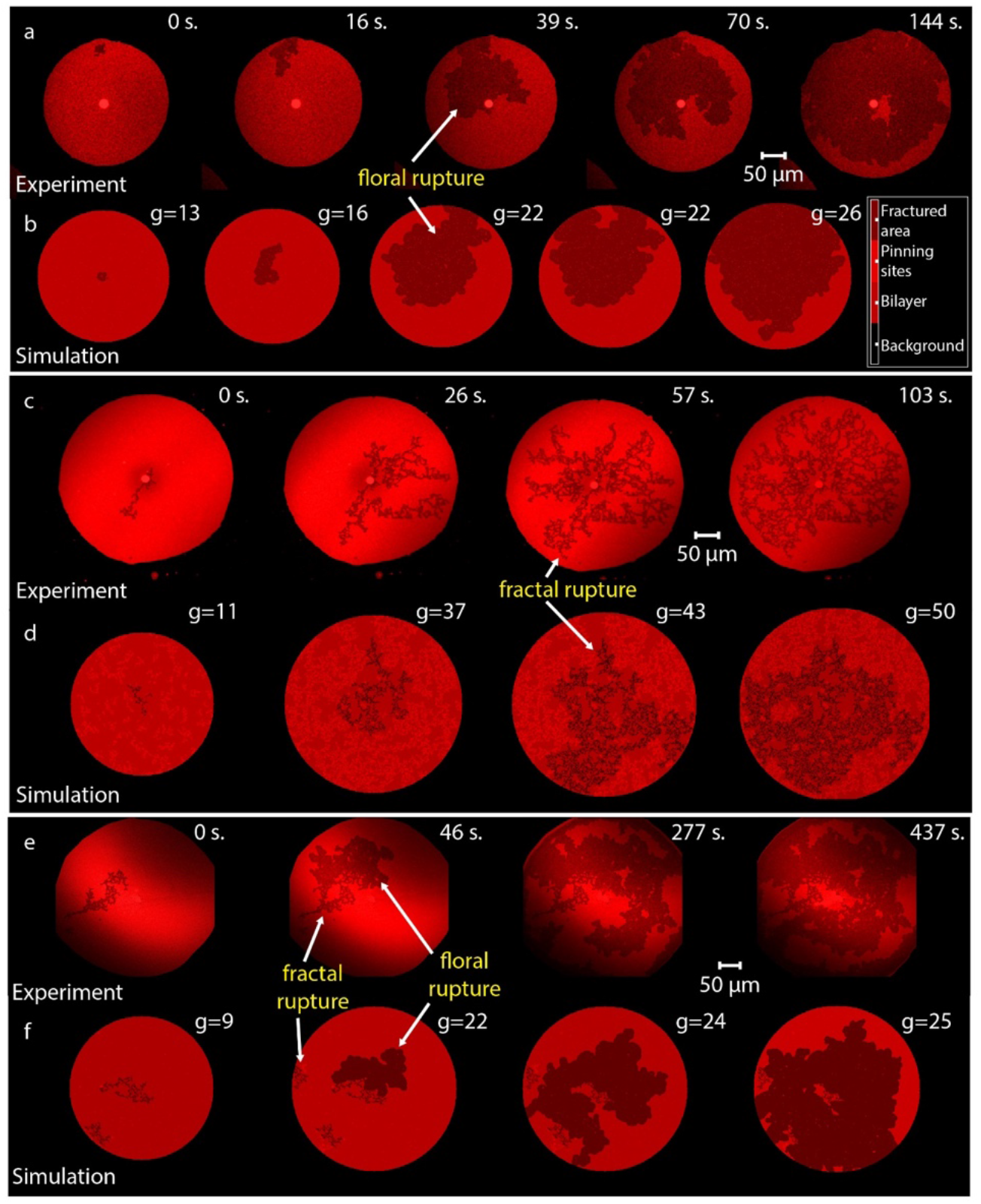
| Experiments and corresponding simulations. The opening and propagation of a floral pore in a circular membrane patch from top view in **(a)** an experiment and **(b)** simulation. The membrane in the experiments is doped with a fluorescence dye which makes it visible. In the simulations different colors represent from the most to less intense: the pinning sites, the bilayer (distal membrane), fractured area (the proximal membrane) and the background (surface). Fracturing of a circular membrane patch in **(c)** an experiment and **(d)** in simulations, from top view. The fractures appear in fractal morphology therefore are called the fractal ruptures. The membranes displaying the fractal and floral ruptures simultaneously **(e)** in an experiment and **(f)** in simulations. *g* represents the CA generation number.

Besides the peculiar characteristic morphology of the fractal pores, the rupture dynamics also differ from the floral ruptures as well as from the most commonly observed circular pores in biomembranes: the fractal ruptures appear intermittently where, in between each occurrence, there is an eventless period. This waiting time corresponds to the time required for the membrane tension to build up again due to ongoing adhesion. When a pore opens, the membrane relaxes and, for the next rupture to appear, the membrane tension needs to increase again to reach to critical point of lysis. The CA model we have established can successfully anticipate this behavior through the reduced tension *T_r_* defined in (3) (Fig. 5). The reduced tension is the expression of membrane relaxation on new offshoot clusters and is governed by the avalanche parameter *AP*. As *AP* is increased, the waiting time between fractal ruptures increases. This behavior in a typical simulation is shown for several *AP* values in Fig. 5d, where the percentage of fractured lipid cells is plotted against the CA generation. Jumps in these plots correspond to the breaking of a cluster. We see that number of jumps tends to decrease and the number of generations between jumps tends to increase as *AP* increases. These plots show good agreement with similar plots of saltatory behavior observed experimentally in^1, 3^.

**Figure 5.**
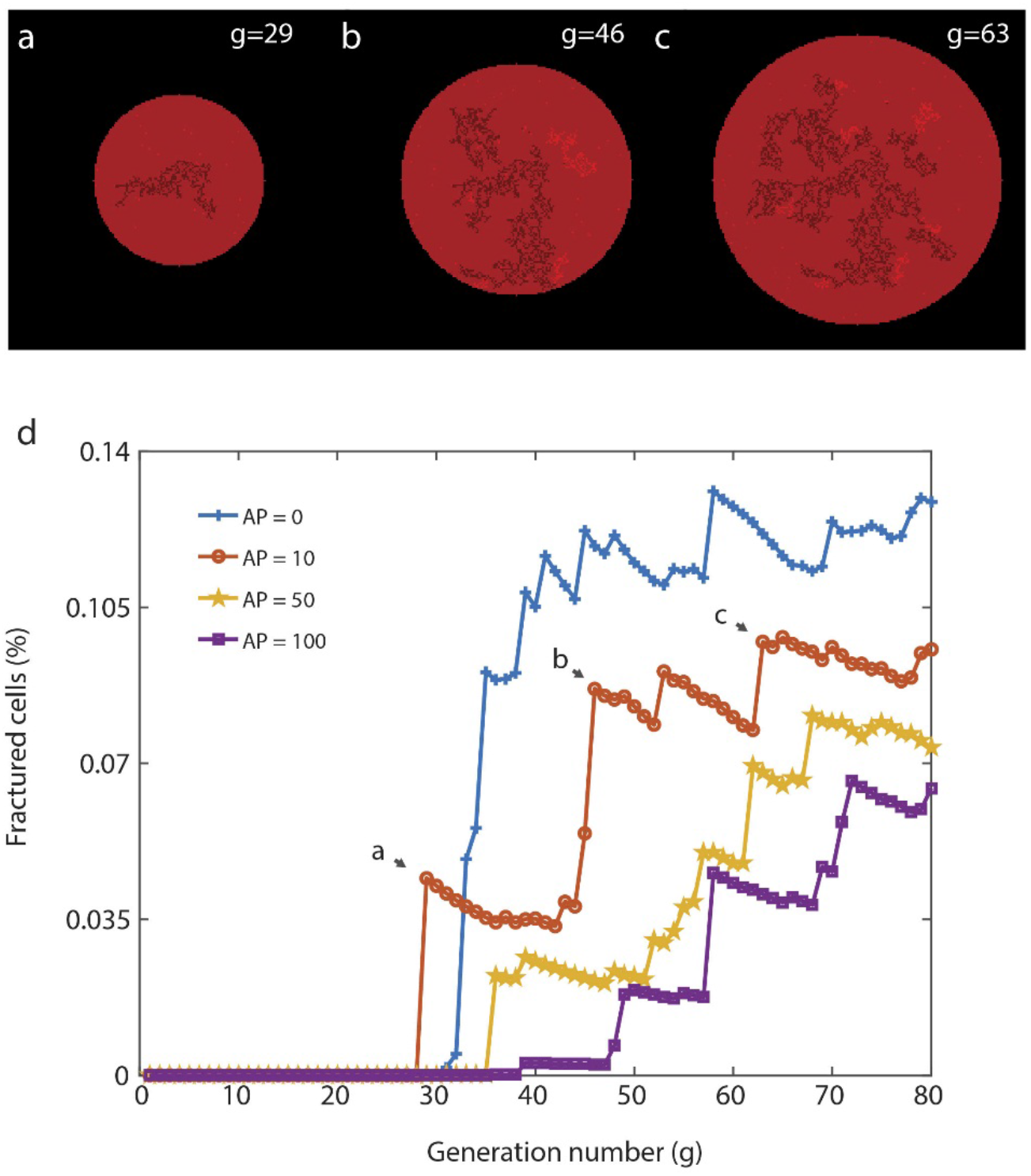
|Intermittency in fractal rupturing. **(a-c)** Simulation snapshots sequentially depicting saltatory fractal rupturing corresponding to an avalanche parameter (AP) of 10. Fractured clusters form new offshoot pinned clusters, which eventually break and form additional cluster. **(d)** Plot showing the percentage of lipid cells fractured in the CA as a function of generation number, demonstrating the effect of the avalanche parameter on the intermittency of fractal rupture. Jumps correspond to a newly fractured cluster. For AP = 0, the delay between cluster ruptures is minimal. As AP is increased, the delay between cluster ruptures is more pronounced.

## Conclusion

In this study, we have proposed a novel cellular automaton that can successfully capture the varied fracture morphology and dynamics of lipid membrane rupture using a small number of parameters and simple rules. Dilute pins with circular ruptures lead to floral rupture morphologies, which spread continuously across the membrane. Cluster pins formed through a percolation process lead to fractal ruptures. Fractal ruptures are punctuated by periods of relaxation, which cause the ruptures to spread intermittently.

It is important to note that the non-trivial ruptures do not occur in extreme conditions but at moderate tensile stress, with similar values to those observed in living cells^28, 29^. Such fractures are therefore anticipated to occur *in vivo*, although the findings are relatively new and currently very little is known regarding the formation of fractures in living organisms. An exception is the earlier observation of fractal pores reported in cultured Chinese Hamster Ovary cells’ plasma membranes^1^. It is high likely that such ruptures continuously occur in tissue as a result of cell-to-cell adhesion or due to pinning of the plasma membrane of the cells to the underlying support: cytoskeleton^30^. Understanding the rules and, ultimately, the physics that underpin these morphologies can help researchers establish the governing factors for the formation of the membrane ruptures thus membrane damage and repair.

## Acknowledgements

GR and IG acknowledge The Research Council of Norway (Forskningsrådet) Project Grant 274433, UiO: Life Sciences Convergence Environment, the Swedish Research Council (Vetenskapsrådet) Project Grant 2015–04561; start-up funding provided by the Centre for Molecular Medicine Norway & Faculty of Mathematics and Natural Sciences at the University of Oslo. MT and AG acknowledge the support of the Kuehler Undergraduate Research Grant and start-up funding provided by the School of Engineering at Santa Clara University.

